# Oral and intranasal Ad5 SARS-CoV-2 vaccines decrease disease and viral transmission in a golden hamster model

**DOI:** 10.1101/2021.10.03.462919

**Authors:** Stephanie N. Langel, Susan Johnson, Clarissa I. Martinez, Sarah N. Tedjakusuma, Nadine Peinovich, Emery G. Dora, Philip J. Kuehl, Hammad Irshad, Edward G. Barrett, Adam Werts, Sean N Tucker

**Author notes:** Correspondence should be addressed to: Stephanie N. Langel, Duke University, Medical Center, 27 Parmer Way, 27703 Durham, NC, USA. and Sean N. Tucker, Vaxart, 170 Harbor Way, South San Francisco, CA, 94080. USA. Contributed equally (Stephanie N. Langel, Susan Johnson).

## Abstract

Transmission-blocking strategies that slow the spread of SARS-CoV-2 and protect against COVID-19 are needed. We have developed a shelf-stable, orally-delivered Ad5-vectored SARS-CoV-2 vaccine candidate that expresses the spike protein. Here we demonstrated that oral and intranasal SARS-CoV-2 vaccination of this candidate protected against disease in index hamsters, and decreased aerosol transmission to unvaccinated, naïve hamsters. We confirmed that mucosally-vaccinated hamsters had robust antibody responses. We then induced a post-vaccination infection by inoculating vaccinated index hamsters with SARS-CoV-2. Oral and IN-vaccinated hamsters had decreased viral RNA and infectious virus in the nose and lungs and experienced less lung pathology compared to mock-vaccinated hamsters post challenge. Naive hamsters exposed in a unidirectional air flow chamber to mucosally-vaccinated, SARS-CoV-2-infected hamsters had lower nasal swab viral RNA and exhibited less clinical symptoms of disease than control animals. Our data demonstrate that oral immunization is a viable strategy to decrease SARS-CoV-2 disease and aerosol transmission.

## Main

The injectable SARS-CoV-2 vaccines currently approved for use are capable of protecting vaccinees from symptomatic infection, hospitalization and death from COVID-19^1-3^. However, they do not completely prevent infection, as evidenced by multiple reports of infections in vaccinated individuals^4,5^. Indeed, mRNA vaccinated individuals infected with the B.1.617 (delta) variant can shed viral RNA and infectious titers and potentially spread SARS-CoV-2 to others^6,7^. Considering most of the world is unimmunized, including children below 12, the possibility that a vaccinated individual with a post-vaccination infection can spread SARS-CoV-2 to unimmunized family or community members poses a public health risk. There would be a substantial benefit to develop vaccines that protect against disease *and* reduce SARS-CoV-2 transmission from vaccinated to unvaccinated individuals.

Since the mucosal surface of the upper respiratory tract (URT) is the initial site of SARS-CoV-2 replication and primary site of infection^8^, interventions that induce robust mucosal immune responses may have the greatest impact on reduction of SARS-CoV-2 transmission. We have created a replication-defective, shelf-stable oral adenoviral vector vaccine candidate expressing the spike (S) protein from SARS-CoV-2 (r-Ad-S) that is designed to induce both systemic and mucosal immunity. Importantly, immune activation via the intestine may represent an important organ for oral immunization as antibody secreting plasmablasts and plasma cells can traffic from the gut to the nose, trachea, and lung^9,10^. Prior work in a human influenza challenge study with the same platform has shown an ability to limit viral RNA shedding of influenza virus^11^, highlighting the utility of this vaccination strategy for respiratory viruses.

To study the potential impact of oral vaccination on transmission to naïve individuals, we used a hamster infection and aerosol transmission system. We vaccinated index hamsters with oral r-Ad-S, using intranasal (IN) r-Ad-S as a control for mucosal stimulation, intramuscular spike protein (IM S) as a control for systemic stimulation, and oral PBS as a mock control. We then infected animals, via IN delivery, with a high titer of SARS-CoV-2 in order to replicate a post-vaccination infection. One-day post viral challenge, index hamsters were placed upstream of vaccine-naïve hamsters in a chamber that allowed aerosol movement but not direct contact or fomite transmission. We demonstrated that post-vaccination, the oral and IN r-Ad-S groups had higher serum IgG and IgA, as well as higher bronchoalveolar lavage (BAL) IgA compared to control animals. This corresponded to decreased SARS-CoV-2 RNA and infectious virus in the URT and decreased weight loss and lung pathology. Importantly, oral and IN r-Ad-S vaccination decreased aerosol transmission of SARS-CoV-2 and reduced disease indicators such as lung inflammation and weight loss in unvaccinated naïve animals, despite the presence of substantial viral RNA in nasal swabs of index immunized animals. These data demonstrate that oral r-Ad-S immunization resulted in reduced disease and decreased SARS-CoV-2 transmission in a hamster model.

## RESULTS

### Oral and IN r-Ad-S vaccination induced robust systemic and mucosal antibody responses

Index hamsters were immunized at weeks 0 and 4 with oral r-Ad-S, IN r-Ad-S (mucosal positive control), IM S protein or mock (oral PBS) prior to SARS-CoV-2 challenge at week 7 (Figure 1A). To determine immunogenicity of these vaccines, serum was collected at weeks 0, 3 and 6 post immunization. BAL was collected upon necropsy (day 5 post-inoculation) (Figure 1A). Oral and IN r-Ad-S vaccinated groups had significantly higher S-specific IgG antibody titers in serum at week 3 compared to mock-dosed hamsters; this was not true in IM S vaccinated hamsters (Figure 1B). Using a surrogate virus neutralizing test (sVNT), serum from IN r-Ad-S hamsters had greater ability to block binding of SARS-CoV-2 S to ACE-2 after the booster vaccination (week 6) compared to serum from mock-vaccinated hamsters (Figure 1C).

**Fig.1:**
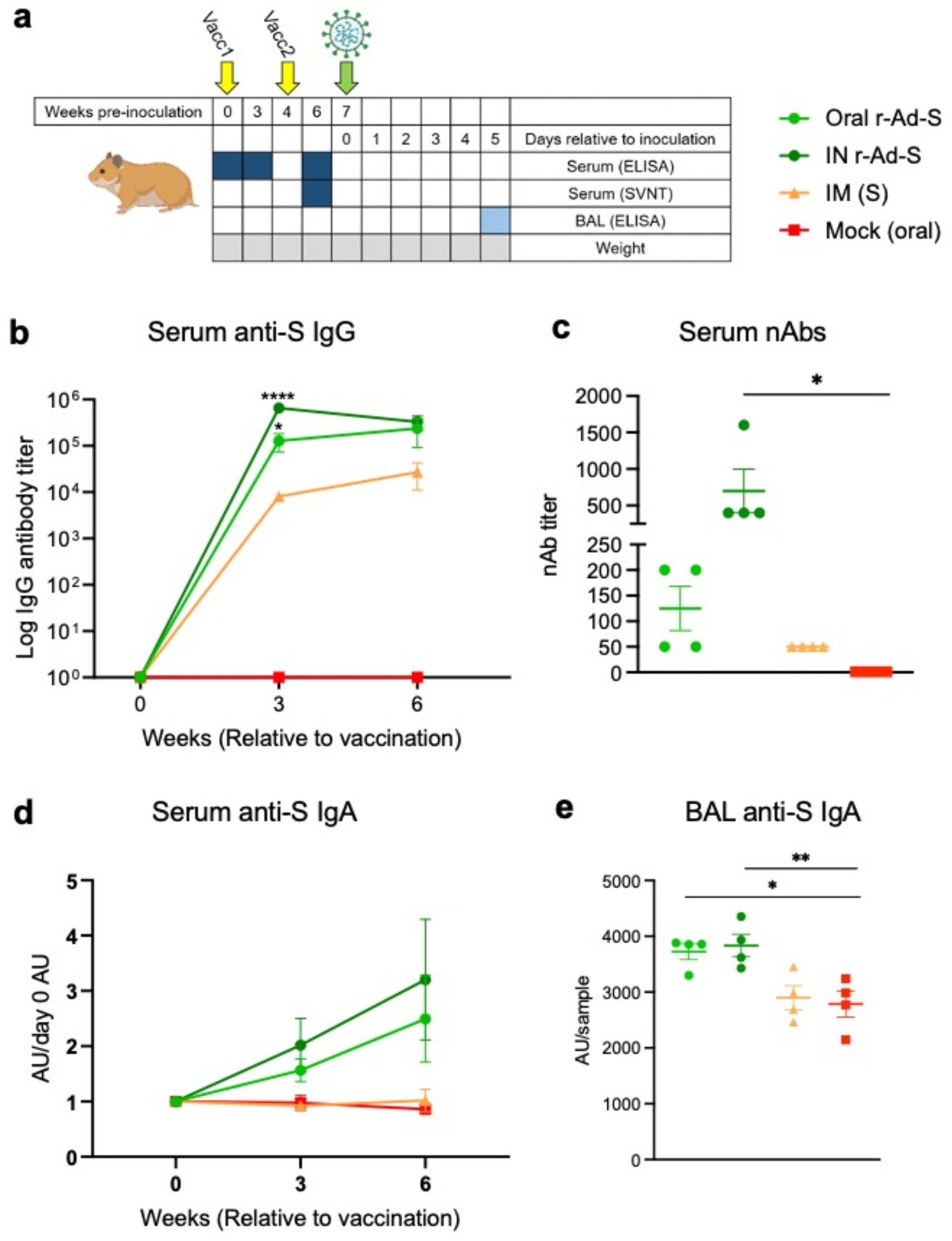
Oral and intranasal r-Ad-S vaccination induced robust IgG and IgA antibodies in golden hamsters. **a-d**, Index hamsters were immunized with oral r-Ad-S, intranasal (IN) r-Ad-S, intramuscular (IM) spike (S) or mock (oral) (*n* = 4 per group) and inoculated with SARS-CoV-2 seven weeks later. **a**, Experimental design schematic. **b**, Serum anti-S IgG antibody titers were measured at weeks 0, 3 and 6 post immunization via ELISA. **c**, SARS-CoV-2 neutralizing antibody (nAb) titers were measured in serum at week 6. **d**, Serum anti-S IgA antibody titers were measured (AU/sample) at weeks 0, 3 and 6 post immunization via Meso scale discovery (MSD) and normalized to day 0 AU values. **e**, anti-S IgA antibodies were measured in bronchoalveolar lavage (BAL) fluid via MSD. Data were analyzed by a one-way ANOVA and Dunnett’s multiple comparisons. Error bars represent the standard error of the mean (SEM). * *P* <0.05, ** *P* <0.01, **** *P* < 0.0001.

Serum anti-S IgA antibodies increased post-oral and IN r-Ad-S vaccination but not in IM S or mock groups (Figure 1D). As expected from our oral immunization platform, oral r-Ad-S vaccinated hamsters demonstrated similar anti-S IgA levels in BAL compared to the IN r-Ad-S positive control group, suggesting similar stimulation of mucosal immunity (Figure 1E). There were no differences in serum or BAL IgA responses between IM S and mock-vaccinated hamsters, suggesting that IgA was not induced by systemic immunization with the IM S protein. These data demonstrate that oral r-Ad-S and IN r-Ad-S vaccinated hamsters generated robust systemic and mucosal humoral immunity.

### Oral and IN r-Ad-S vaccination accelerated SARS-CoV-2 viral RNA clearance and protected against disease in hamsters

In index animals, SARS-CoV-2 RNA from nasal swabs at 3 and 5 days post-inoculation was lower in oral and IN r-Ad-S-vaccinated animals compared to mock-vaccinated animals; this was not true for IM-vaccinated animals (Figure 2A). Nasal swabs of index animals immunized with IN r-Ad-S had significantly lower TCID_50_ values compared to mock animals on day 1 (Figure 2B), but all groups still had detectable infectious virus. Oral and IN r-Ad-S-vaccinated index animals had lower viral RNA loads in the lungs at terminal collection (day 5) when compared to mock-vaccinated animals, which was not true for IM S-vaccinated animals (Figure 2C). All vaccinees had significantly reduced lung infectious viral loads, with all appearing below the limit of detection at day 5 (Figure 2D). These data demonstrated that oral and IN r-Ad-S vaccination decreased SARS-CoV-2 replication post infection, and accelerated viral clearance.

**Fig.2:**
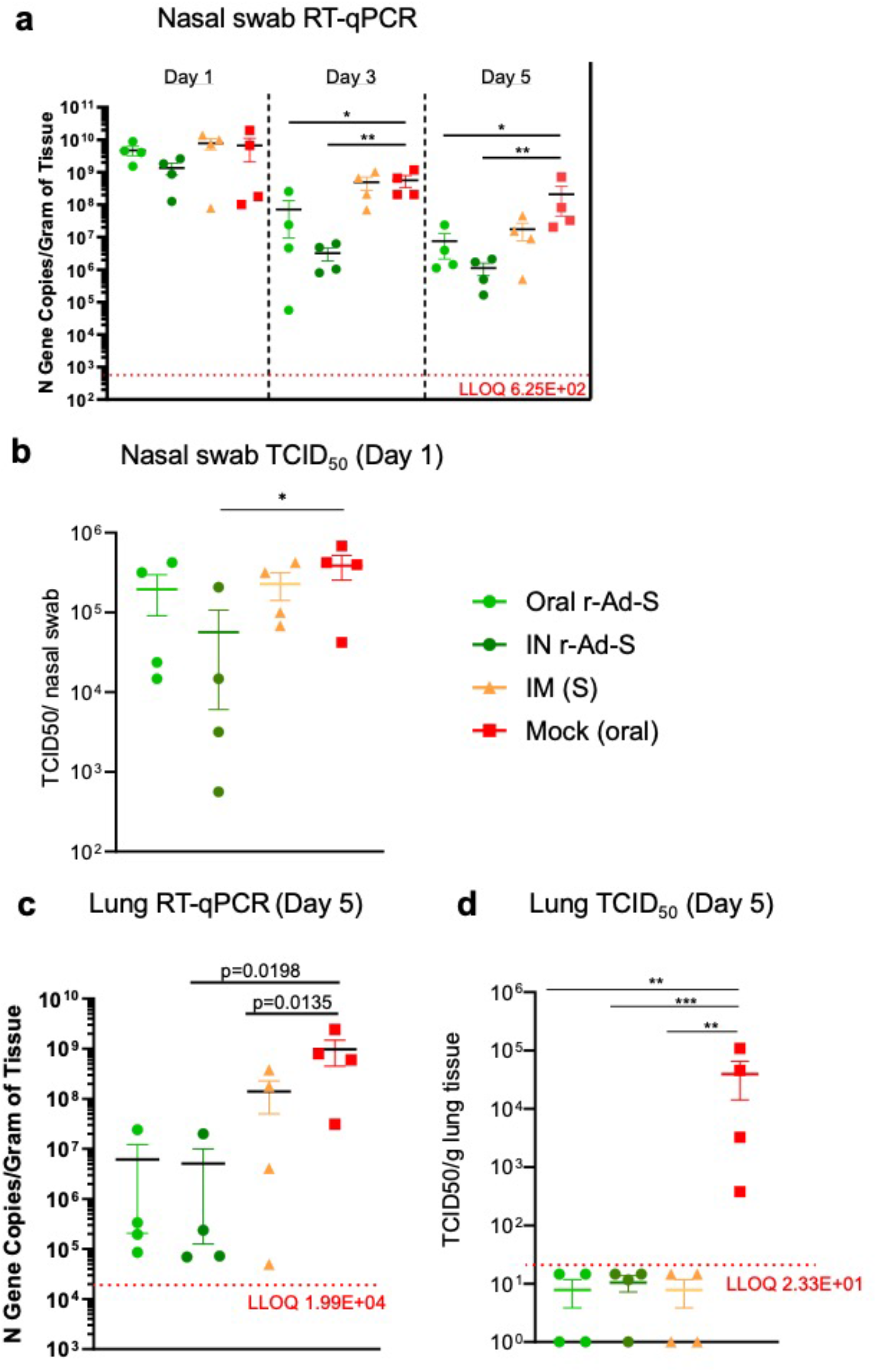
Oral and intranasal r-Ad-S vaccination decreased SARS-CoV-2 RNA and infectious virus in nasal swabs and lungs. **a**, Nasal swabs were collected on days 1, 3 and 5 in index hamsters and viral RNA loads in these samples were determined by quantitative reverse transcription PCR (qRT-PCR) of the N gene. **b**, Nasal swabs were collected on day 1 and infectious virus titers were determined by TCID_50_. **c**, Lung tissue was collected at necropsy (day 5) and RNA was isolated for SARS-CoV-2 detection by qRT-PCR of the N gene and **d**, infectious viral titers by TCID_50_. The dotted line represents Lower Limit of Quantification (LLOQ), with data below the limit of detection plotted at ½ LLOQ. Data were analyzed by a one-way ANOVA and Dunnett’s multiple comparisons. Error bars represent the SEM. * *P* <0.05, ** *P* <0.01, *** *P* < 0.001.

During vaccination but prior to SARS-CoV-2 inoculation, all animals were gaining weight at a similar rate (Extended data Fig. 1). After SARS-CoV-2 inoculation, we observed significantly greater weight loss in unvaccinated hamsters (AUC = 767 ± 2.5) compared to uninfected hamsters (AUC = 809.6 ± 4.0) (Figure 3A), a characteristic of disease in this model. Oral and IN r-Ad-S vaccinated animals lost less weight by the termination of the study compared to mock-vaccinated animals (Figure 3B). To quantify pulmonary inflammation, lung weights were measured, and lungs were scored for gross pathology. In index animals, lung weights (normalized to total body weight) (Figure 3C) and average gross pathology scores (Figure 3D) were decreased in both oral and IN r-Ad-S-vaccinated groups when compared to mock-treated hamsters.

**Fig.3:**
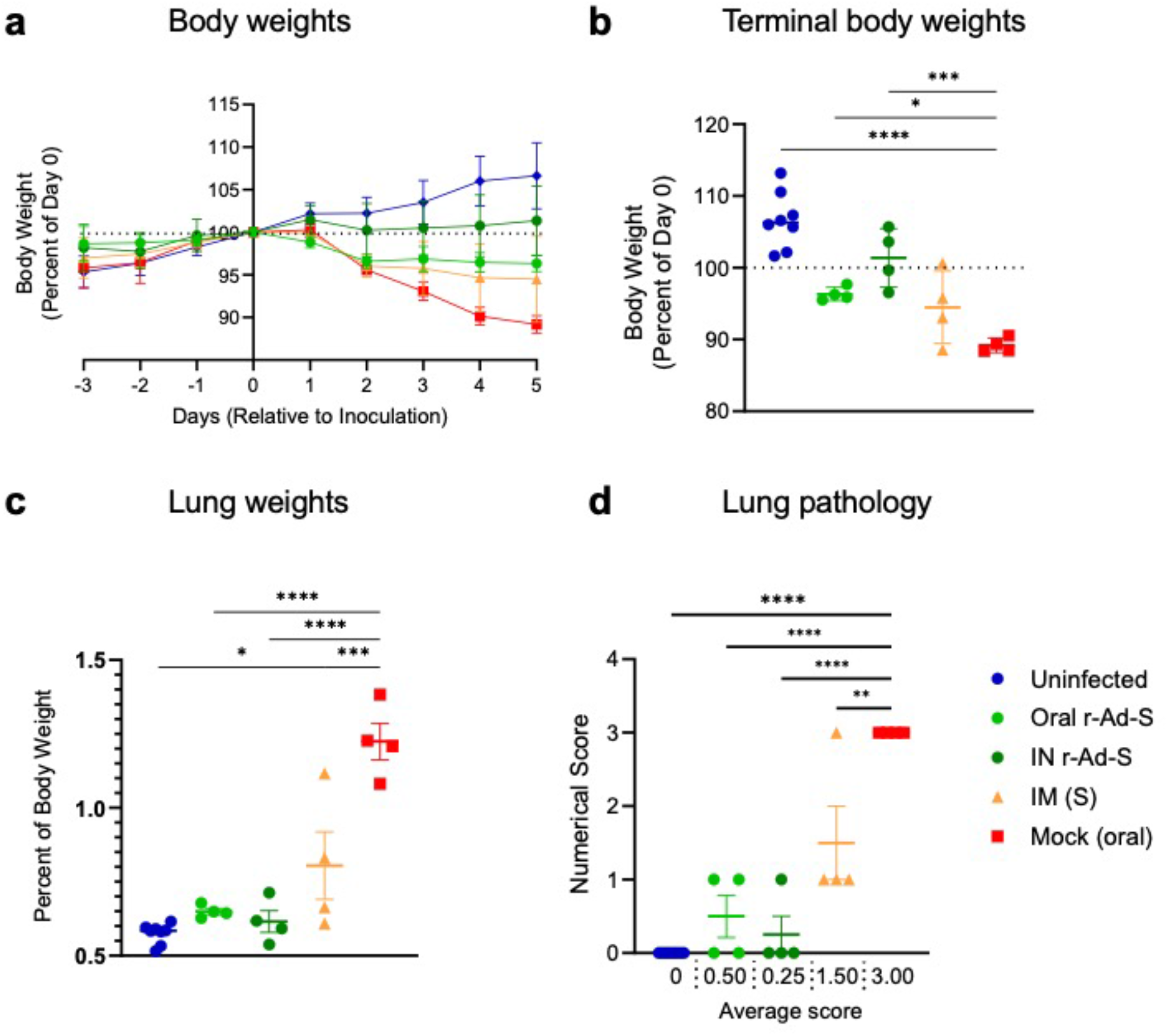
Oral and intranasal r-Ad-S vaccination reduced disease indicators including body weight, lung weight and lung pathology scores in index hamsters. **a**, Daily weight change and **b**, terminal body weights were determined by the percent of day 0 (relative to SARS-CoV-2 inoculation). **c**, terminal lung weights and **d**, lung pathology scores were determined. Severity grade for red discoloration of the lung was based on a 0 to 4 scale indicating percent of whole lung affected: none (no grade), minimal (1), mild (2), moderate (3), marked (4) correlating to 0, 1-25, 26-50, 51-75, and 76-100% affected, respectively. Data were analyzed by a one-way ANOVA and Tukey’s multiple comparisons. Error bars represent the SEM. \**P* <0.05, ** *P* <0.01, *** *P* < 0.001, **** P<0.0001.

### Oral and IN r-Ad-S vaccination limited SARS-CoV-2 transmission to unvaccinated, naïve hamsters leading to decreased clinical indicators of disease

As a test of transmissibility, unvaccinated naïve hamsters were exposed to vaccinated, SARS-CoV-2-infected index animals at a 1:4 ratio of index:naive, where each vaccine exposure group has 4 index animals and 16 vaccine naïve animals. Exposure was performed by putting an index animal in a chamber (index chamber) connected to a second chamber containing the four naïve animals (naïve chamber), separated by a 5 inch connecting chamber (Extended data Fig. 2). Screens at either end of the connecting chamber prevented direct contact. Air was circulated in the index chamber by fan, and was pulled into the naïve chamber by vacuum. Exposure of naïve hamsters to aerosols produced by index hamsters was performed for 8 hours prior to moving each hamster to an individual cage. On day 1 post infection the amount of viral RNA loads in all index animals was above 7.8×10^7^ gene copies per swab (Figure 2A).

In naïve hamsters exposed to the oral and IN r-Ad-S-immunized groups, SARS-CoV-2 RNA was significantly lower on days 1 and 3 compared to mock-immunized animals (Figure 4A). The number of vaccine naïve animals with nasal swab viral RNA levels above a threshold of 1×10^5^ gene copies was also determined. On day 1 post exposure of the index and naïve animals, there were 3 naïve hamsters (3/16) exposed to the oral r-Ad-S index group that were above the threshold compared to eleven mock exposed hamsters (11/16) (p=0.011, Fisher’s exact test) (Figure 4A, Extended data Table 1). The naïve hamsters exposed to IN r-Ad-S index hamsters had 0 (0/16) animals above the threshold which was not significantly different than oral, but significantly lower than naïve animals exposed to aerosol from mock-vaccination animals (p=0.22, p<0.0001 by Fisher’s exact test respectively) (Figure 4A, Extended data Table 1). On day 3, in the oral exposed group, 11 (11/16) had nasal swab viral RNA levels above the threshold compared to 16 (16/16) for the naïve exposed group (p=0.043 by Fisher’s exact test) (Figure 4A, Extended data Table 1). The naïve animals exposed to IN index animals had 6 (6/16) which was not significantly different than oral (p=0.16 by Fisher’s exact test) (Figure 4A, Extended data Table 1). These data demonstrate decreased SARS-CoV-2 transmission from oral and IN r-Ad-S**-**vaccinated index to naïve animals. On day 5, nasal viral RNA did not differ between groups. However, lung viral RNA and infectious virus were significantly lower in IN r-Ad-S vaccinated compared to unvaccinated hamsters (Figure 4B-D).

**Fig.4:**
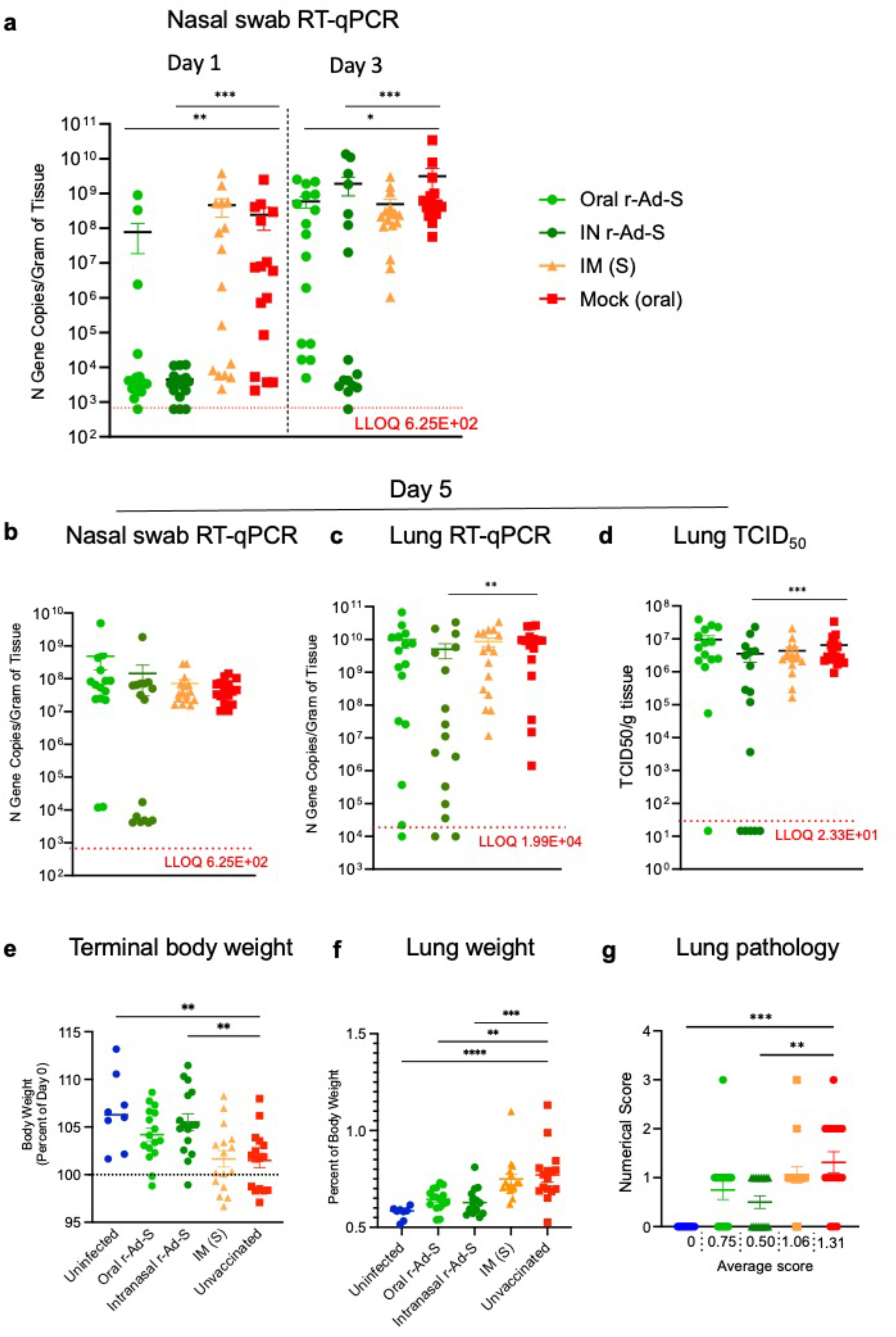
Oral and intranasal SARS-CoV-2 vaccination decreased SARS-CoV-2 transmission and clinical indicators of disease. **a**, Nasal swabs were collected in naïve animals on days 1, 3 and **b**, 5 after exposure to index, infected hamsters in aerosol chamber. Viral RNA loads in these samples were determined by quantitative reverse transcription PCR (qRT-PCR) of the N gene. **c**, Lung tissue was collected at necropsy (day 5) and RNA was isolated for SARS-CoV-2 detection by qRT-PCR of the N gene and **d**, infectious viral titers were determined by TCID_50_. **a-d**, The dotted line represents LOD, with data below the limit of detection plotted at ½ LOD. Data were analyzed by a one-way ANOVA and Dunnett’s multiple comparisons. **e**, terminal body weights were determined by the percent of day 0 (relative to SARS-CoV-2 inoculation). **f**, Terminal lung weights and **g**, lung pathology scores were determined. Severity grade for red discoloration of the lung was based on a 0 to 4 scale indicating percent of whole lung affected: none (no grade), minimal (1), mild (2), moderate (3), marked (4) correlating to 0, 1-25, 26-50, 51-75, and 76-100% affected, respectively. **e-g**, Data were analyzed by a one-way ANOVA and Tukey’s multiple comparisons. **a-g**, Error bars represent the SEM. \**P* <0.05, ** *P* <0.01, *** *P* < 0.001, **** P<0.0001.

While oral and IN r-Ad-S vaccination didn’t completely prevent transmission, the vaccination likely reduced the effective dose reaching the naïve animals. In a prior virus titering experiment, we demonstrated animals lost more weight and had increased pulmonary inflammation when they received higher titers of SARS-CoV-2 inoculum, suggesting that the severity of disease acquired is dependent on the original infectious dose (Extended data Fig. 3). Similar dose-dependent disease severity findings have been shown by others in hamsters, mice, ferrets and nonhuman primates^12-16^. Reduction in effective dose may have led to a greater proportion of naïve animals exposed to oral r-Ad-S (14/16) and IN r-Ad-S (15/16) vaccinated hamsters gaining weight by the end of the study (Figure 4E). This is compared to naïve animals exposed to the IM (S) vaccinated (8/16) and unvaccinated (10/16) hamsters (Figure 4E) where fewer naïve animals increased in size. Additionally, oral and IN r-Ad-S-vaccinated hamsters had lower lung weights (Figure 4F) and pathology (Figure 4G) scores compared to unvaccinated hamsters. These data demonstrate that oral and IN r-Ad-S-vaccinated animals with a post-vaccination SARS-CoV-2 infection transmit less infectious virus via aerosol to unvaccinated, naïve hamsters than infected but unvaccinated (or IM vaccinated) animals, and that this difference in transmission resulted in less severe clinical outcomes.

## DISCUSSION

SARS-CoV-2 emerged in late 2019 and quickly spread around the globe, leading to hundreds of millions of cases and over 4 million deaths. Despite reports of decreased viral RNA shedding from mRNA-vaccinated individuals compared to unvaccinated individuals^7,17^, recent evidence demonstrates that vaccinated individuals can get infected with SARS-CoV-2 and shed infectious virus, which can lead to onward transmission. Improved mucosal responses that stimulate local immunity at the site of SARS-CoV-2 replication may have a much greater impact on human transmission. In this study, we show that oral and IN r-Ad-S vaccination decreases disease in vaccinated hamsters and reduces SARS-CoV-2 transmission to unvaccinated naïve hamsters.

To induce post-vaccination infection, we infected index animals with a high dose of SARS-CoV-2, and subsequently exposed naïve animals to these vaccinated, infected index animals in an aerosol exposure chamber. We observed faster clearance of viral RNA in oral and IN r-Ad-S-vaccinated compared to mock-vaccinated hamsters. Additionally, aerosol transmission to naïve animals during the 8-hr exposure window was substantially reduced from oral and IN r-Ad-S-vaccinated hamsters as evidenced by reduced nasal swab viral RNA loads in naïve animals one and three days post transmission exposure. We hypothesize that mucosal antibodies in the URT were able to enhance SARS-CoV-2 clearance in vaccinated animals and limit the infectiousness of transmitted aerosols. Consistent with this hypothesis, anti-S IgA in the BAL of the oral and IN r-Ad-S immunized animals was higher than IM or mock vaccinated animals. Other groups have also tested SARS-CoV-2 transmission in a hamster model. For example, van Doremalen and colleagues determined that IN-vaccinated hamsters were protected from SARS-CoV-2 when placed in the same cage as unvaccinated hamsters inoculated with 1×10^4^ TCID_50_ of virus^18^. These experiments determined whether IN-vaccinated individuals were protected against transmission from unvaccinated individuals. In our study, we tested whether mucosal vaccination reduces the ability of the virus replicating in vaccinated individuals from being transmitted specifically via aerosol to naïve animals. We used a relatively high dose of virus to induce a post-vaccine infection (1×10^5^ TCID_50_) and physically separated animals in the exposure chamber to remove the possibility of contact/fomite exposure as a means of transmission. We show for the first time that mucosal vaccination can reduce SARS-CoV-2 transmission from vaccinated to unvaccinated animals. These data are of high relevance because of its implication in global public health, especially in areas in which a significant percentage of the people are unvaccinated. Additionally, if new variants arise that result in an increase in post-vaccine infections, reducing transmission even in vaccinated populations would contribute to limiting circulating virus. Mucosal vaccination could be considered for implementation not only because they protect the vaccinee, but because they likely have a greater effect on the community as a whole. Reducing SARS-CoV-2 transmission to the unprotected is likely to lead to decreased hospitalization and deaths.

Our study does have limitations. Firstly, we do not measure mucosal T cell responses which are shown to play a role in limiting SARS-CoV-2 infection at mucosal sites^19^. Additionally, the SAR-CoV-2 challenge dose we used was above physiological dose likely to be picked up by an environmental exposure, as evidenced by the high viral RNA load in the nose at 1-day post infection in all index hamsters. It is likely that mucosal immunization would provide greater protection against SARS-CoV-2 transmission when lower doses of challenge virus are used. Our study was meant to be a stringent challenge to clearly identify advantages between vaccine groups. For practical reasons, such as the limitation of the number of aerosol transmission chambers, lower dose levels or evaluation at later time points were not done. Additionally, future work should include challenging mucosally-vaccinated hamsters with the Delta variant and other variants of concern.

An orally-delivered, temperature-stable SARS-CoV-2 vaccine is ideal for global vaccination, where adequate storage and qualified health care providers maybe in short supply. IN delivery has some of the same advantages, but translating IN SARS-CoV-2 vaccination efficacy in humans has proven to be more difficult than in animals^20,21^. Implementing oral vaccine campaigns around the world has been done, as evidenced by the rotavirus and poliovirus vaccination efforts^22,23^. We have previously demonstrated that our SARS-CoV-2 clinical candidate vaccine VXA-COV2-1 generated robust humoral immune responses in mice^24^. Additionally, it was tolerated and immunogenic in a phase I clinical trial where the oral vaccine was delivered to people as tablets (https://clinicaltrials.gov/show/NCT04563702) and protected golden hamsters from SARS-CoV-2 challenge when given by oral gavage (Johnson, 2021, submitted). An additional phase 2 study using the vaccine candidate tested in this study and given as a tablet will begin clinical studies shortly. In summary, the data presented here demonstrate that oral immunization is a viable strategy to decrease SARS-CoV-2 transmission and disease, and should be considered for vaccination efforts that increase global immunity to SARS-CoV-2.

## Acknowledgements

The experimental timeline schematic was created using BioRender (www.biorender.com). This work was supported by the Bill and Melinda Gates Foundation award INV-022595 (S.N.L.). The following reagent was produced under HHSN272201400008C and obtained through BEI Resources, NIAID, NIH: Spike Glycoprotein (Stabilized) from SARS-Related Coronavirus 2, Wuhan-Hu-1 with C-Terminal Histidine Tag, Recombinant from Baculovirus, NR-52308. Thanks to Dr. Anne Moore for reviewing the manuscript.

## Author information

These authors contributed equally: Stephane N. Langel, Susan Johnson

## Affiliations

**Duke Center for Human Systems Immunology, Department of Surgery, Duke University Medical Center, Durham, NC, USA**

Stephanie N. Langel

**Vaxart, San Francisco, CA, USA**

Susan Johnson, Clarissa I. Martinez, Sarah N. Tedjakusuma, Nadine Peinovich, Emery G. Dora, Sean N. Tucker

**Lovelace Biomedical Research Institute**

Philip J. Kuehl, Hammad Irshad, Edward G. Barrett, Adam Werts

## Contributions

S.N.L., S.J., E.G.B., A.W. and S.N.T. conceived, designed and analyzed the experiments. P.J.K. and H.I. designed and constructed the aerosol transmission chamber. H.I. and A.W. carried out the in vivo aerosol transmission experiments. N.P. and E.G.D. created the vaccine material. C.I.M. and S.N.T. performed the immunological experiments. S.N.L., S.J., A.W. and S.N.T wrote the manuscript with input from all co-authors.

## Corresponding authors

Correspondence to Stephanie N. Langel or Sean N. Tucker.

## METHODS

### Vaccine constructs

r-Ad-S is a rAd5 vector containing full-length SARS-CoV-2 S gene under control of the CMV promoter. rAd5 vaccine constructs were created based on the published DNA sequence of SARS-CoV-2 publicly available as Genbank Accession No. MN908947.3. The published amino acid sequences of the SARS-CoV-2 S were used to create recombinant plasmids containing transgenes cloned into the E1 region of Adenovirus Type 5 (rAd5)^25^, using the same vector backbone used in prior clinical trials for oral rAd tablets^11,26^. All vaccines were grown in the Expi293F suspension cell-line (Thermo Fisher Scientific) and purified by CsCl density centrifugation. The S Protein vaccine was provided by BEI Resources, NR-52308. Spike Glycoprotein (Stabilized) from SARS-Related Coronavirus 2, Recombinant from Baculovirus, NR-52308.

### Animals and study design

Male Syrian Hamsters (*Mesocricetus auratus*) approximately 12-14 weeks of age with a weight range of 106-136g were sourced from Charles River Laboratory. Animal work was performed at Lovelace Biomedical, with approval from the Institutional Animal Care and Use Committee. Hamsters were singly housed in filter-topped cage systems and were supplied with a certified diet, filtered municipal water, and dietary and environmental enrichment. The study was powered to compare viral RNA loads in naïve hamsters exposed to vaccinated, SARS-CoV-2 infected index animals between vaccine groups, where beta was set to 0.2, alpha=0.05. Assuming an attack rate of 80% infected in the placebo group and a vaccine efficacy of 70%, an N=15 was calculated with continuity correction^27^. The study was rounded to 16 naïve, 4 index to maintain the 1:4 ratio. The study was not powered to directly compare index groups, but many statistical significances were achieved with N=4.

All r-Ad-S vaccinations were given at a dose of 1e9 IU (1:100 of a human dose^11^). Oral vaccine was delivered by gavage in 300μl of PBS subsequent to delivery of 300μl 7.5% bicarbonate buffer. IN vaccination was delivered in PBS by pipette, 50μl/nostril. The control group received PBS via oral gavage. One group received recombinant SARS-CoV-1 S protein, made in insect cells, by the IM immunization group (BEI, #NR-722). All index animals were challenged by IN inoculation of SARS-CoV-2 at approximately 1E+05 TCID_50_/animal in 300µl volume seven weeks after the initial vaccination. Index animals were then housed for 24 hrs individually before placing in aerosol chamber. Each vaccine/index group had 4 animals, and matched with a corresponding N=16 naïve exposed animals (1:4 ratio of index to naïve, with 1 index animal exposed to 4 naïve in a chamber setup). All animals were sacrificed five-days post inoculation (index) or aerosol chamber exposure (naïve) for terminal assays.

### Transmission chamber

The aerosol transmission chamber was based on previous transmission work^15^ and adapted for hamsters. The chamber consists of multiple subchambers that support unidirectional flow. Starting from left to right in Extended data Fig. 2: 1) chamber for the index hamster, 2) a connector chamber, and 3) a chamber for the naïve hamster(s). All chambers were fitted with access doors with air tight seals and appropriate safety features to ensure animals cannot interact with fans, sampling, flow, etc. Approximate dimensions of the first chamber (1) is 4”x10”x9” (length, width, height). The unidirectional flow (5 L/min) was controlled by regulated house exhaust flow from the 3^rd^ chamber. This drew room air into chamber 1 via a HEPA filter (F; HEPA Vacuum Filter Compatible with Kenmore 86880, EF-2, Panasonic MC-V194H Vacuum Cleaner). Chamber 1 was also fitted with a recirculating fan (D; ANVISION 40mm x 10mm DC 5V USB Brushless Cooling Fan, Dual Ball Bearing, Model YDM4010B05) to ensure homogeneity of the aerosol prior to transitioning into chambers 2 and 3. The connector chamber (chamber 2) (approximate dimensions of 4”x5”x5”), connected chambers 1 and 3 in order to separate the hamsters, but allows for air passage. Chamber 3 (approximately 10”x10”x9”) houses the naïve hamsters. A wire mesh screen with 0.25”x 0.25” holes was placed at each end of chambers 1 and 3 to prevent hamsters moving to another chamber. Additionally, chamber 2 was raised off the ground to prevent feces and/or urine from moving between chambers.

### Assessment of infectious SARS-CoV-2 load in lung homogenate

Lung tissue samples from euthanized hamsters infected with SARS-CoV-2 were collected, weighed and homogenized with beads using a Tissue Lyser (Qiagen). Each sample was serially diluted 10-fold in Dulbecco’s Modified Eagle’s Medium containing 2% fetal bovine serum (FBS) and 1% Pen-Strep solution. VeroE6 cell monolayers at ≥ 90% confluency in 96-well plates were rinsed with PBS. The plates were inoculated with 100µl of each sample dilution in five technical replicates. Negative control wells contained dilution medium only. The plates were incubated at 37°C and 5% CO_2_ for 72 hours. Cytopathic effect was scored after fixing the cell monolayers with 10% formalin and staining with 0.5% crystal violet. Viral load was determined in tissue culture infectious dose 50%/ml of lung homogenate using the Reed and Muench method^28^. Infectious virus titers in infected lungs were expressed as TCID50/gr of tissue.

### SARS-CoV-2 Surrogate Neutralization Assay

Neutralizing antibodies were measured using a SARS-CoV-2 Surrogate Virus Neutralization Test Kit (Genscript). Hamster sera was diluted from 1:20 to 1:500 incubated at a 1:1 ratio with HRP conjugated SARS-CoV-2 RBD protein for 30 min at 37°C. Following incubation, 100 μl was added to hACE2 pre-coated plates and incubated for 15 min at 37 °C. Plates were washed 4x with 260μl per well of supplied wash solution, followed by the addition of 100 μl per well of supplied TMB solution. Plates were developed for 15 min at RT in the dark before development was stopped with 50 μl per well of supplied stop solution. OD was measured at 450 nm with a Spectra Max M2 microplate reader.

### IgG ELISA

Purified SARS-CoV-2 S1 protein (GenScript) in carbonate buffer, pH 9.4, (Thermo Scientific) as coated onto microtiter plates (Maxisorp, Nunc) at 1 μg/ml and incubated overnight at 4°C before blocking with 100 μl of PBS-0.05% Tween (PBST) + 1% BSA for 1 hr. Serum samples were serially diluted in PBST. After a 2-hr incubation at room temperature (RT) the plates were washed 3x with PBST, followed by the addition of 100μl per well of 1:3000 goat anti-hamster IgG-HRP (Thermo Fisher) in PBST + 1% BSA. Plates were incubated at RT for 1 hr before washing 3x with PBST before the addition of 50μl per well of TMB substrate (Rockland). The plates were developed for 10 minutes, then stopped with 50μl per well of 2M sulfuric acid. Optical densities (OD) were measured at 450nm with a Spectra Max M2 microplate reader.

### IgA MSD

S1 protein was biotinylated according to manufacturer’s instructions (EZ-link, Thermofisher), and was conjugated to a U-Plex MSD linker (Mesoscale Diagnostics). The linked S1 protein was coated on 2-spot U-plex 96 well plates (Mesoscale Diagnostics) for a final concentration of 66nM per well over night. The next day the plates were blocked with PBS-T for 1 hour prior to the addition of hamster serum or BAL. Samples from each individual hamster acquired on D0, D28 and D55 were quantified on the same plate. The serum samples were diluted to 1:200 and the BAL samples were added to the plate neat. Following 2-hr sample incubation, the plate was washed and SULFO-TAG (MSD) anti-hamster IgA (Brookwood Biomedical) detection antibody was added. The plate was washed and 1X MSD read buffer was added and each plate was analyzed using MSD QuickPlex.

### SARS-CoV-2 Challenge

All animals were challenged by IN inoculation of 1×10^5^ TCID_50_/animal SARS-CoV-2 (isolate USA-WA1/2020) at 100µl/nostril eight weeks post initial vaccination. SARS-CoV-2, isolate USA-WA1/2020, was sourced from BEI Resources and propagated in Vero E6 African Green Monkey kidney cells (BEI, catalog #N596) at the University of Texas Medical Branch (UTMB). Virus was stored in a biosafety level 3 compliant facility.

### Detection of SARS-CoV-2 in lung homogenates by qRT-PCR

Lung samples were weighed and homogenized with beads using a Tissue Lyser (Qiagen) in 1 ml of TRI reagent, before RNA was isolated and purified from tissue samples using the Direct-Zol 96-RNA kit (Zymo Research). Copies of SARS-CoV-2 N were measured by qRT-PCR TaqMan Fast Virus 1-step assay (Applied Biosystems). SARS-CoV-2 specific primers and probes from the 2019-nCoV RUO Assay kit (Integrated DNA Technologies) were used: (L Primer:TTACAAACATTGGCCGCAAA; R primer: GCGCGACATTCCGAAGAA; probe:6FAM-ACAATTTGCCCCCAGCGCTTCAG-BHQ-1). Reactions were carried out on a Stratagene MX3005P or BioRad CFX384 Touch instrument according to the manufacturer’s specifications. A semi-logarithmic standard curve of synthesized SARS-CoV-2 N gene RNA (LBRI) was obtained by plotting the Ct values against the logarithm of cDNA concentration and used to calculate SARS-CoV-2 N gene in copies per gram of tissue.

### Gross pathology scoring

Gross necropsy observations of the lung will be recorded in Provantis using consistent descriptive terminology to document location(s), size, shape, color, consistency, and number. Gross observations will include a severity grade for red discoloration of the lung (likely to be associated with pneumonia) based on a 0 to 4 scale indicating percent of whole lung affected: none (no grade), minimal (1), mild (2), moderate (3), marked (4) correlating to 0, 1-25, 26-50, 51-75, and 76-100% affected, respectively.

