## Supplemental for "Oral and intranasal Ad5 SARS-CoV-2 vaccines decrease disease and viral transmission in a golden hamster model"

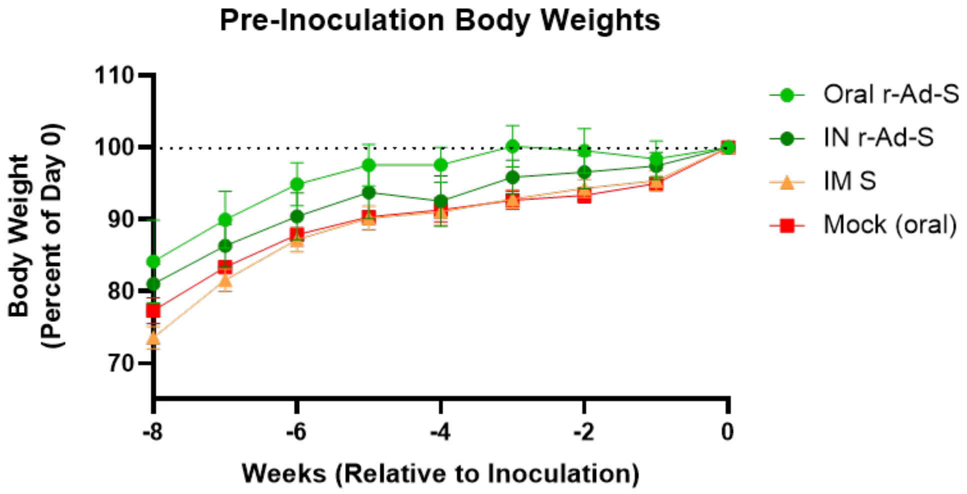


**Extended Data Fig. 1. All animals gained weight on a similar trajectory prior to SARS-CoV-2 inoculation.** Animals were weighed weekly and data was graphed as a percent of day 0 (SARS-CoV-2 inoculation day) in oral r-Ad-S, intranasal (IN) r-Ad-S, intramuscular (IM) spike (S) and mock (oral) vaccinated golden hamsters. Error bars represent the SEM.

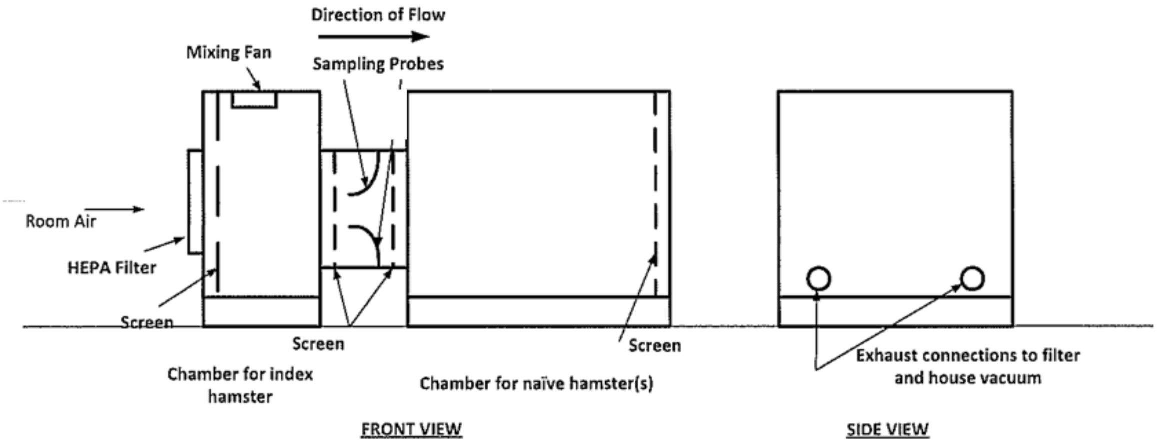


**Extended Data Fig. 2. Aerosol transmission chamber.**

**Extended Table 1. Outcome table of animals at or above $1\times10^5$ gene copies**

**Day 1. Gene Copies $1\times10^5$ or Above**

| Group | Positive | Negative | p versus oral | Significant |
|---|---|---|---|---|
| Oral rAd-S | 3 | 13 | NA | NA |
| Intranasal rAd-S | 0 | 16 | 0.22 | No |
| Spike protein vaccine | 10 | 6 | 0.029 | Yes |
| Mock | 11 | 5 | 0.011 | Yes |

**Day 3. Gene Copies $1\times10^5$ or Above**

| Group | Positive | Negative | p versus oral | Significant |
|---|---|---|---|---|
| Oral rAd-S | 11 | 5 | NA | NA |
| Intranasal rAd-S | 6 | 11 | 0.16 | No |
| Spike protein vaccine | 16 | 0 | 0.043 | Yes |
| Mock | 16 | 0 | 0.043 | Yes |

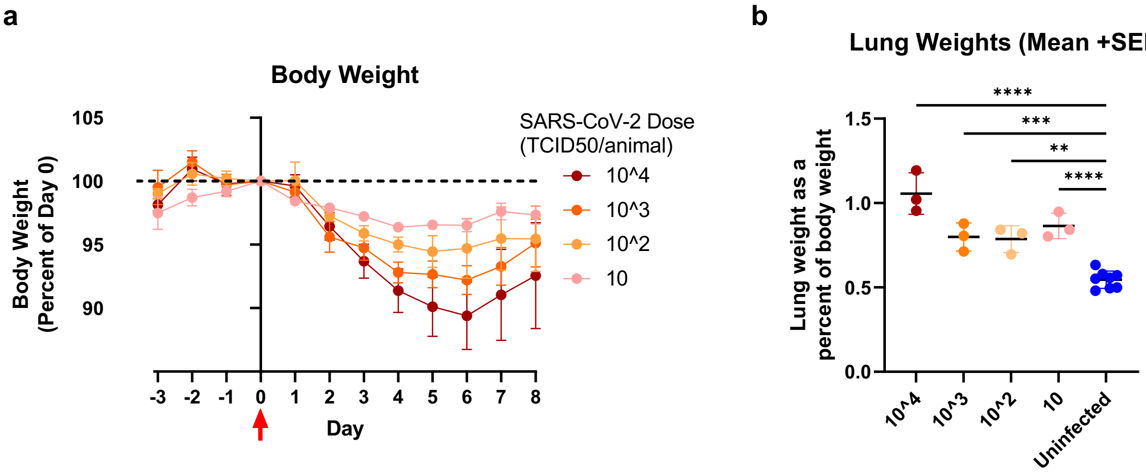


**Extended Data Fig. 3. Disease severity varies directly in accordance to the viral titer at inoculation.**

**a,** Hamsters were inoculated with 1×10, 1×$10^2$, 1×$10^3$, or 1×$10^4$ $TCID_{50}$ of SARS-CoV-2 and body weights were determined for 8 days. **b,** Lung weight as a percent of body weight was determined at day 8 post inoculation. Error bars represent the SEM. ***P* <.01, ****P* <.001, *****P* <.0001,
